# Phylotastic: improving access to tree-of-life knowledge with flexible, on-the-fly delivery of trees

**DOI:** 10.1101/419143

**Authors:** Van D. Nguyen, Thanh H. Nguyen, Abu Saleh Md. Tayeen, H. Dail Laughinghouse, Luna L. Sánchez-Reyes, Enrico Pontelli, Dmitry Mozzherin, Brian O’Meara, Arlin Stoltzfus

**Affiliations:** Department of Computer Science, New Mexico State University, Box 30001, MSC CS, Las Cruces, 88003, New Mexico, USA; Institute for Bioscience and Biotechnology Research, 9600 Gudelsky Drive, Rockville, 20850, MD, USA; Ft. Lauderdale Research and Education Center, University of Florida/IFAS, 3205 College Avenue, Davie, 33314, FL, USA; Department of Ecology and Evolutionary Biology, University of Tennessee, 569 Dabney Hall, Knoxville, 37996, TN, USA; Illinois Natural History Survey, Species File Group, University of Illinois, 1816 South Oak St., Champaign, 61820, Illinois, USA; Office of Data and Informatics, NIST, 100 Bureau Drive, Gaithersburg, 20899, MD, USA

## Abstract

(1) A comprehensive phylogeny of species, i.e., a tree of life, has potential uses in a variety of contexts in research and education. This potential is limited if accessing the tree of life requires special knowledge, complex software, or long periods of training.

(2) The Phylotastic project aims to use web-services technologies to lower the barrier for accessing phylogenetic knowledge, making it as easy to get a phylogeny of species as it is to get online driving directions. In prior work, we designed an open system of web services to validate and manage species names, find phylogeny resources, extract subtrees matching a user’s species list, calibrate them, and mash them up with images and information from online resources.

(3) Here we report a publicly accessible system for on-the-fly delivery of phylogenetic knowledge, developed with user feedback on what types of functionality are considered useful by researchers and educators. The system currently consists of a web portal that implements 3 types of workflows to obtain species phylogenies (scaled by geologic time and decorated with thumbnail images); 19 underlying web services accessible via a common registry; and toolbox code in R and Python so that others can create applications that leverage these services. These resources cover most of the use-cases identified in our analysis of user needs.

(4) The Phylotastic system, accessible via http://www.phylotastic.org, provides a unique resource to access the current state of phylogenetic knowledge, useful for a variety of cases in which a tree extracted quickly from online resources (as distinct from a tree custom-made from character data) is sufficient, as it is for many casual uses of trees identified here.

## 1 Introduction

As Cracraft, et al. [3] suggest, a tree of life “provides a comparative and predictive framework for all fundamental and applied biology.” Yet, obtaining a suitable phylogeny is difficult. Inferring a phylogeny from raw data following best practices is a formidable task, and phylogeny experts typically do not employ practices that make their phylogenies discoverable, accessible, and re-useable [4, 11, 23]. Indeed, most trees are generated for a narrow and specific purpose. Nevertheless, literature surveys show that trees are re-used in research, typically by subtree extraction from large species trees [23]. That is, from a tree with broad coverage, a subtree is extracted covering only the paths of descent for a subset of descendant nodes, typically tip-nodes representing species of interest.

The prospects for disseminating tree-of-life knowledge via subtree extraction has increased greatly with the OpenTree project [7], which currently provides a “synthetic tree” with over 2 million species constructed by a supertree method from 819 source trees [20]. OpenTree’s database contains these and many more source trees awaiting curation, all available via a programmable web-services interface, as are trees from TreeBASE [18] and Phylomatic [24]. Other types of relevant information accessible via web services include taxonomies [2, 17], occurrence records from GBIF (Global Biodiversity Information Facility) [5] or iNaturalist, and images and other information (size, habitat, etc) from the Encyclopedia of Life (EOL) [16].

Based on such resources, one may envision integrative discovery tools with a wide variety of uses, operating within an ecosystem of web services, e.g., a tool to get an annotated phylogeny for all of the birds identified in a particular country. The conception of such a system—– a “Phylotastic” system focused on improving the discoverability and accessibility of phylogenetic knowledge—– along with a proof-of-concept implementation, has been previously described [22].

Here we describe a robust and accessible implementation of a Phylotastic system, featuring a web portal that implements several workflows to specify a desired output tree based on various inputs (documents, web sites, user lists, taxon searches), a set of 31 web services, a web-services registry, and two function libraries (in R and Python) to facilitate development of software that takes advantage of these services. We explain the functionality of these tools relative to a set of use-cases based on interviews with prospective users, and compare them to other tools currently available.

## Analysis and design

### Use cases

The original concept for a Phylotastic system [22] emanated from research scientists with expertise in phylogenetics and informatics. To ensure broader appeal, we developed a Phylotastic web portal and then used it to obtain feedback (via correspondence as well as in-person interviews) from a broader range of potential users, including educators at multiple levels, as well as researchers who were not experts in informatics or phylogenetics. Based on this information, we prioritized the following use-cases.

#### Generate a tree from a specified list of taxa

Provide a tree for a user-supplied list of species or higher taxa. For example, a botanist wants to demonstrate the relationships of 10 named genera of interest within the family Asteraceae.

#### Generate a tree of N species sampled from a named taxon

Given a named taxon and a number N, provide a tree with N species chosen in some way, e.g., at random, by popularity, or by maximal diversity (taxonomic or phylogenetic). For instance, a user wants to illustrate the diversity of rodents using a tree with no more than 25 species.

#### Generate a phylo-guide from an electronic resource

Create a tree with images and links to species information from a web page or document that includes taxonomic names, such as a scientific paper, an online encyclopedia (e.g., EOL, Wikispecies), or a document listing the species found in a park, zoo, region, or collection. For example, an educator working from a zoo’s species list wants to create a guide that students can use to discover information during a field trip; a researcher evaluating a paper wishes to see all of the named taxa in a phylogenetic context; a web developer wants to build a plugin that will show the phylogeny of species implicated in any web page.

#### Contextualize high-level relationships

Given the problem of illustrating a simple general relationship of a set of taxa, such as {plants, animals, fungi}, generate a tree using familiar species that illustrate the relationship, possibly including species from other (unspecified) taxa for context. For example, pick three species per taxon to show the relationships of birds, turtles, crocodilians, lizards, snakes, and mammals.

#### Integrate data or metadata with phylogeny

Given the set of species implicated by any method described above, return a tree and an associated data table integrating information or resources of interest, including images, links to information (EOL or wikipedia), or data on features such as toxicity, pathogenicity, availability of fossil data, medicinal value, conservation status, size, biogeography, or habitat.

Currently the system we describe partially covers these cases (see Discussion). Some desirable types of data are not available systematically (e.g., medicinal value, pathogenicity), and some types of operations are not implemented in available algorithms (e.g., choosing a set of species based on both popularity and diversity).

### Implicated workflows and operations

The kinds of operations identified previously [22] include (1) rectifying names by matching input strings with qualified taxonomic names; (2) finding available trees with coverage of user-identified taxa; (3) extracting a subtree; (4) assigning branch lengths by some means, e.g., fossil scaling; (5) adding data or metadata for species or higher taxa; (6) rendering a tree graphically. The expanded set of use cases above implicates a slightly larger set of operations, including (7) extracting taxonomic names embedded in other text, and (8)sampling members of a taxonomic group by some criteria.

### Web services registry and support for automated planning

Further requirements are induced, not directly by the use cases above, but by the desire for an open system that can be maintained as a community resource [22]. Web services, which provide access to data and operations from any point on the internet via standard protocols, are an obvious choice for a decentralized community system. In many cases, there is a one-to-one correspondence between operations above and web services that exist or could exist. For example, there are multiple Taxonomic Name Resolution Services (TNRSs) that take an input list and match it against a taxonomy [2, 15].

Given a rich set of web services, it will be possible to develop diverse *client* applications that use the system, each drawing on some combination of web services. A *web services registry* is used to make the available set of web services discoverable. The registry provides information on what each service does, and how to access it. Sponsors of participating web services may register their services, providing a description that makes the services discoverable and accessible.

Finally, the system we describe here is designed to support automated fault-tolerant knowledge discovery. This leads to a requirement for a machine-readable registry that supports methods of automated planning and workflow composition. The use of this system in automated planning has been described elsewhere [12, 13], and is not directly relevant to the functionality that we describe here.

## Implementation

### Service registry

We implemented a *Web Service Registry (WSR)* as a software application to register, store, discover, and execute web services. The WSR is a Web-based application that includes an application layer and a database layer used to store the description of the available web-services, including where to locate the service and how to invoke it. To support automated knowledge discovery, web services are described in a machine-readable way using WSDL, the *Web Service Description Language (WSDL)*. In the application layer, the WSR provides a Content Management System (CMS) based on Drupal technology, ^1^ where the managed contents are web service objects.

### Services

In general, Phylotastic services are designed for synchronous operation, to support workflows carried out on-the-fly, with results being returned to users in real time. One exception is a set of services to manage persistent lists, so that a list created by a client in a session may be accessed in a later session, or by a different client. For instance, a mobile birding application could create a list of bird species, authenticated with a Google account, that is accessed later using the Phylotastic web portal.

Currently there are 31 phylotastic services that fall into the categories below. Some of these services are thin wrappers around external services, while others were developed for this project.

#### Taxonomic name resolution

These services are used for matching input strings to formal taxonomic names. For example, *GNR_TNRS_wrapper* accepts scientific names as input and returns resolved names with identifiers from known taxonomy sources such as Catalog of Life or NCBI using the Global Names Resolver ^2^. *NCBI_common_name* takes a list of common names (vernacular names) and suggests scientific names based on the NCBI taxonomy ^3^.

#### Scraping names

These services extract scientific names embedded in the text of electronic resources including web pages, PDFs, Microsoft Office documents, and even images. For example, *GNRD_wrapper_URL* accepts the URL of a web resource (which may be an HTML page or an electronic file) and identifies scientific names based on services provided by the Global Names Recognition and Discovery (GNRD) project ^4^.

#### Taxon sampling

The services in this category take the name of a higher taxon and return a list of species chosen by some criterion. For example, the *Taxon_genome_species* takes a named taxon and returns a list of species within that taxon for which there is a genome sequence available from NCBI.

#### List management

The services in this category are used to publish, access, remove or update lists of species. For example, *Add_new_list* service allows any valid user (users with a gmail address) to insert a new list of species into the list server.

#### Taxon information and images

These services fetch information on named species or higher taxa. For example, *EOL_Habitat_Conservation* takes a list of species and returns the habitat and conservation status, using the EOL traits bank. *Image_url_species* takes a list of species names as input and obtains image URLs (and corresponding license information), using the service provided by EOL ^1^.

#### Phylogenetic tree retrieval

The services in this category return phylogenetic trees. For example, *OToL_wrapper_Tree* accepts a list of taxa and returns a tree. This service first uses the Open Tree of Life’s (OToL) *match_names* method API to get the Open Tree Taxonomy (OTT) identifiers of the input taxa, and then uses the *induced_subtree* method to retrieve the Phylogenetic tree.

#### Tree scaling

These services can be used for fitting a tree to a geologic time scale. For example, *Datelife_scale_tree* accepts a Phylogenetic tree (in Newick format) and produces a tree with branch lengths using the Datelife R package ^2^.

#### Tree comparison

*Compare_trees* accepts two trees (in Newick format) and returns true if the trees are the same. This service, currently used only in automated tests, is implemented via DendroPy ^3^.

### Libraries in R and Python

To facilitate the development of client software, we have developed toolbox code in R and Python, installable packages that provides access to Phylotastic services using functions written in the native language.

The rphylotastic package covers nearly all of the categories of services described above. Functions call a phylotastic web service and translate the results with the package jsonlite [14], a single list of species names from any higher taxon of interest to the user can be obtained with the command `rphylotastic::taxon_get_species(“<name>”)`.

The package structure is standard, including function documentation, a manual and a vignette. Function names follow ropensci’s style (https://github.com/ropensci/onboarding/blob/master/packaging_guide.md). Documentation and manual were generated with roxygen2 R package [19]. To evaluate rphylotastic’s package performance and robustness, a set of unit tests were designed and implemented using the R package testthat [19]. Around 58 % of the package code is currently covered by this test suite. ^4^

The phylotastic python package^5^ has methods to access almost all of the services from each category described above. The main module of the python package is *phylotastic_services* module which have different sub-modules implementing different services.

To access a service, a user needs to import the main module and invoke the corresponding method of the service. For example, to get a Phylogenetic tree from Open Tree of Life source, a user needs to call the method *get_tree_OpenTree* with a list of taxa as input.

The documentation for the python package was generated with Sphinx^1^ python documentation generator. To test the functional correctness of the python package submodules, a set of unit tests were implemented using Python Unit Testing Framework and deployed in Travis-CI^2^.

### Web portal

The web portal is written in Ruby using Rails, a model-view-controller (MVC) framework for rapid development of robust web applications. The portal takes advantage of PostgreSQL for database management; Paperclip for managing file attachments; Twitter-Bootstrap, JQuery, and FontAwesome for front-end development; Devise for authentication management; Wicked PDF for PDF generation; Capybara and Minitest for automated testing; Docker and Kubernetes for containerization and deployment. The test suite covers model tests, controller tests, and interactive tests (simulated in Poltergeist, which mimics user interactions).

The portal supports three types of workflows shown in Fig. 1: extract names from an electronic resource (e.g., document file, web page) that may contain text other than names; upload a formatted list of names; and sample species from a named taxon. To reduce barriers to prospective users, no login is required to access these workflows, e.g., a user may extract species names from a web page specified by its URL, create a named list, use this to create a named tree, and customize the tree visualization. These features are accessible anonymously in a session-dependent manner. When the user logs in, lists and trees associated with a session will be migrated to the user’s account and will be accessible from other devices.

**FIgure 1:**
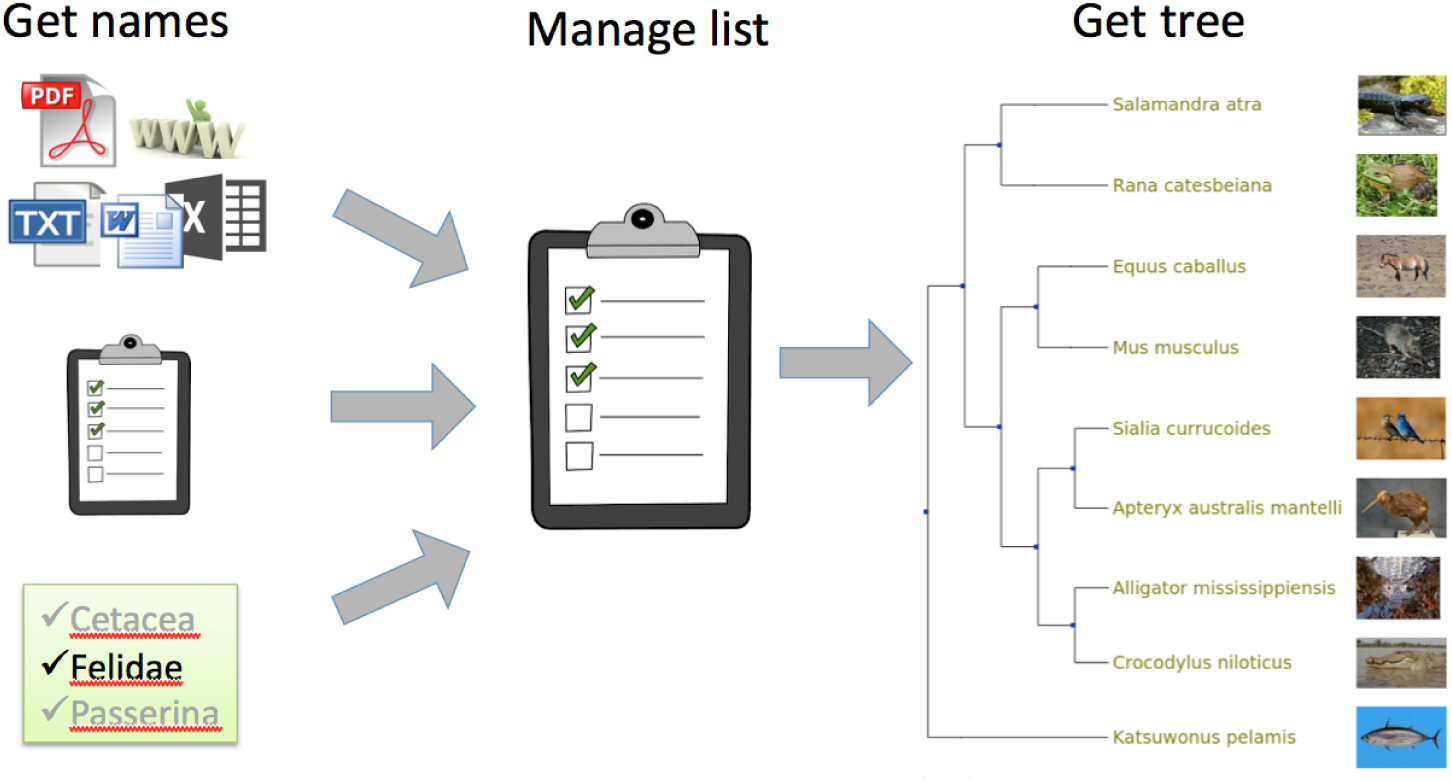
Phylotastic portal workflows.

## Availability

Code, applications and services may be discovered and accessed via the project’s web home at http://www.phylotastic.org. The web portal is accessible via http://portal.phylotastic.org, and the registry is at http://registry.phylotastic.org. Source code is available under an open-source license from the GitHub Phylotastic organization (http://www.github.com/phylotastic). At present, all resources are accessible without restriction; usage restrictions on web services may be imposed in the future if necessary to ensure that the services are broadly usable. The stable version of rphylotastic can be installed from R directly from the CRAN repository (https://cran.r-project.org/package=rphylotastic) using *install.packages(pkgs = “rphylotastic”)*. Development versions are available from GitHub repository (https://github.com/phylotastic/rphylotastic) and can be installed using *devtools::install_github(“phylotastic/rphylotastic”)*. The Python library can be installed following the instructions at https://github.com/phylotastic/phylotastic_py.

## Discussion

### Support for use-cases

The Phylotastic web portal illustrates some of the potential of the Phylotastic system, and is designed to serve as a useful resource for scientists, educators and students. The general workflow for use of the portal is (1) initiate a list of taxa; (2) manage the list; (3) get a tree and render it graphically. The rendering may be customized, and either the tree or the tree-rendering can be downloaded. There are various ways to designate a list of species

- use a public list already available in the portal
- upload a list, as a text file with one name per line, or as a DwC-A archive
- scrape names embedded in an uploaded file (PDF, doc, txt, xls, image)
- scrape names from a resource URL (HTML or other resource)
- from a named taxon, choose
  - a random sample of *N* species
  - species with known genomes (via NCBI services)
  - species with occurrence records in a given location (via iNaturalist)

From this description of capabilities it will be apparent that the portal provides the capacity to generate a tree from a user-supplied list of species or higher taxa (use case 1), to generate a tree of N species from a chosen taxon (use case 2), and to generate a tree for names embedded in an electronic resource such as a document or web site (use case 3).

Providing phylogenetic context to illustrate a relationship (use case 4) can be done in stages by creating a separate list of sampled species for each taxon using the portal, downloading the lists with the portal, concatenating them in a text editor, then uploading the full list for tree retrieval and display using the portal.

The portal can integrate some kinds of metadata or data with a phylogeny (use case 5), specifically EOL links and thumbnail images. One way to integrate other data is to begin with an electronic resource combining the desired type of data for the desired list of species, and then using that as an input to the portal, e.g., scraping the wikipedia page entitled “List of poisonous plants” yields a list of 185 names, for which list the portal will return a tree with 153 tips. If the user has a spreadsheet with data for a set of species, the names can be scraped out of the spreadsheet to obtain a tree, and the matrix and tree can be combined using a web tool to generate visualizations of trees with data, such as EvolView [6] or IToL [9].

### Comparison with other resources

The resources of this project can be considered broadly as a way of making tree-of-life knowledge accessible, through both a set of tools to support the development of software (client applications including scripts), and a multi-purpose interactive tool, the Phylotastic web portal. Other resources available to satisfy a user’s interest in phylogenetic knowledge provide for (1) computing a trees from comparative data (e.g., sequence alignments);(2) discovery and retrieval of pre-computed trees (e.g., from TreeBASE); (3) interactive browsing of a tree of life; or (4) subtree extraction from supertrees and other large trees.

Currently no resources are comparable to the Phylotastic project in the breadth of support for the use-cases listed above. ToLWeb [10], OneZoom [21], IToL [9] and the OpenTree web portal all allow interactive browsing of a tree of life, but they do not support extracting relationships for an arbitrary subset of species (except via a web-services API in the case of OpenTree). Many tools provide support for obtaining gene or protein sequences, aligning them, and inferring a phylogeny. Even the most highly automated of these systems requires judgment and advanced skills, e.g., SUPERSMART [1] is a complex system that must be installed and run by someone proficient in command-driven tools. A better alternative for most users would be a system that provides computing services, such as CIPRES, or more conveniently, Phylogeny.fr.

The most obviously comparable resources to the web portal are Phylomatic [24] and TimeTree [8], both of which helped to inspire the Phylotastic project. Both are interactive web tools that support extraction of subtrees. The web portal offers a greater variety of ways to specify a set of taxa. The main focus of TimeTree is on providing a time-scale for evolutionary divergence based on published chronograms (time-calibrated trees). The Phylotastic portal also provides scaled trees using DateLife (an open version of Time-Tree), although its store of chronograms is smaller.

## Author contributions

VDN and ASMD implemented the portal, and contributed to design and testing, along with AS and HDL; AS and HDL analyzed requirements for the portal and web services; LLSR and BO designed, implemented and tested the DateLife web portal, DateLife web services, and the RPhylotastic package; ASMD and THN implemented web services, and contributed to design and testing along with AS; ASMD designed and implemented the phylotastic_py package; DM designed and implemented improvements to taxonomic name resolution services; AS, BO and EP conceived of the project and oversaw all aspects of design and testing; AS drafted the manuscript, which was completed with the help of the other authors.

## Acknowledgements

This work was supported by funding from the US National Science Foundation (award 1458572, “Collaborative Research: ABI Development: An open infrastructure to disseminate phylogenetic knowledge”). The identification of any specific commercial products is for the purpose of specifying a protocol, and does not imply a recommendation or endorsement by the National Institute of Standards and Technology. We thank Yan Wong, James Rosindell, Karen Cranston, Jonathan Rees, Jaime Huerta-Cepas, Brian Sidlauskas, Jim Allman, Ramona Walls, Mark Holder, Daisie Huang, Cyndy Parr, Katja Schulz, and Son Tran, for useful comments, discussions, and technical advice. We thank many others who contributed indirectly to this project through their participation in two NESCent hackathons with a Phylotastic theme.

## Footnotes

1 https://www.drupal.org/

2 http://resolver.globalnames.org/

3 https://www.ncbi.nlm.nih.gov/taxonomy

4 http://gnrd.globalnames.org/

1 http://eol.org/api/

2 http://datelife.org/

3 http://dendropy.org/

4 https://codecov.io/gh/phylotastic/rphylotastic/tree/master/R

5 https://github.com/phylotastic/phylotastic_py

1 http://www.sphinx-doc.org/en/1.5/index.html

2 https://travis-ci.org/

